# Both Simulation and Sequencing Data Reveal Multiple SARS-CoV-2 Variants Coinfection in COVID-19 Pandemic

**DOI:** 10.1101/2021.09.06.459196

**Authors:** Yinhu Li, Yiqi Jiang, Zhengtu Li, Yonghan Yu, Jiaxing Chen, Wenlong Jia, Yen Kaow Ng, Feng Ye, Bairong Shen, Shuai Cheng Li

**Affiliations:** Department of Computer Science, City University of Hong Kong, Hong Kong 999077, China; State Key Laboratory of Respiratory Disease, National Clinical Research Center for Respiratory Disease, Guangzhou Institute of Respiratory Health, the First Affiliated Hospital of Guangzhou Medical University, Guangzhou 510120, China; Department of Computer Science, Hong Kong Baptist University, Hong Kong 999077, China; Department of Computer Science, Faculty of Information and Communication Technology, Universiti Tunku Abdul Rahman, Kajang 43000, Malaysia; Institutes for Systems Genetics, Frontiers Science Center for Disease-related Molecular Network, West China Hospital, Sichuan University, Chengdu 610041, China

**Keywords:** SARS-CoV-2 variant coinfection, Viral transmission simulation, Coinfection index, Heterozygous SNPs

## Abstract

SARS-CoV-2 is a single-stranded RNA betacoronavirus with a high mutation rate. The rapidly emerged SARS-CoV-2 variants could increase the transmissibility, aggravate the severity, and even fade the vaccine protection. Although the coinfections of SARS-CoV-2 with other respiratory pathogens have been reported, whether multiple SARS-CoV-2 variants coinfection exists remains controversial. This study collected 12,986 and 4,113 SARS-CoV-2 genomes from the GISAID database on May 11, 2020 (GISAID20May11) and April 1, 2021 (GISAID21Apr1), respectively. With the single-nucleotide variants (SNV) and network clique analysis, we constructed the single-nucleotide polymorphism (SNP) coexistence networks and noted the SNP number of the maximal clique as the coinfection index. The coinfection indices of GISAID20May11 and GISAID21Apr1 datasets were 16 and 34, respectively. Simulating the transmission routes and the mutation accumulations, we discovered the linear relationship between the coinfection index and the coinfected variant number. Based on the linear relationship, we deduced that the COVID-19 cases in the GISAID20May11 and GISAID21Apr1 datasets were coinfected with 2.20 and 3.42 SARS-CoV-2 variants on average. Additionally, we performed Nanopore sequencing on 42 COVID-19 patients to explore the virus mutational characteristics. We found the heterozygous SNPs in 41 COVID-19 cases, which support the coinfection of SARS-CoV-2 variants and challenge the accuracy of phylogenetic analysis. In conclusion, our findings reported the coinfection of SARS-CoV-2 variants in COVID-19 patients, demonstrated the increased coinfected variants number in the epidemic, and provided clues for the prolonged viral shedding and severe symptoms in some cases.

## Introduction

Severe acute respiratory syndrome coronavirus 2 (SARS-CoV-2) has infected more than 176.5 million persons, with more than 3.8 million deaths at the time of preparing this manuscript [1, 2]. The virus is an enveloped and single-stranded RNA betacoronavirus of 30k base-pairs, which belongs to the family Coronaviridae [1]. Since the year 2000, we have witnessed and experienced three highly widespread pathogenic coronaviruses in human populations, and the other two are severe acute respiratory syndrome (SARS)-CoV in 2002-2003, and Middle East Respiratory Syndrome (MERS)-CoV in 2012 [3]. All three viruses can lead to acute respiratory distress syndrome (ARDS) in the human hosts, which may cause pulmonary fibrosis and lead to permanent lung function reduction or death [4]. Although with lower mortality rates than SARS-CoV and MERS-CoV, SARS-CoV-2 could invade host cells by binding to the ACE2 on the host cell surface and cause rapid spread among people [5].

To address the challenges, researchers conducted various studies to explore the genomic sequences of SARS-CoV-2 [6-8]. Qianqian Li *et al*. have analyzed 13,406 spike sequences of SARS-COV-2 variants in the GISAID database and divided the SARS-CoV-2 variants into seven evolutionary groups using neutralizing monoclonal antibodies [6]. Correspondingly, the Centers for Disease Control and Prevention also reported the new emerged SARS-CoV-2 variants that circulating globally, including B.1.1.7 lineage in the United Kingdom, B.1.351 lineage in Nelson Mandela Bay and South Africa, P.1 lineage in Japan and Brazil, B.1.429 lineage in the United States, *etc*. [9]. From Pengfei Wang *et al*.’s study, we learned that the extensive mutations in the spike protein of B.1.1.7 and B.1.351 variants could enhance their resistance to the neutralization by convalescent and post-vaccination sera. These reports enforce the notion that the newly emerged SARS-CoV-2 variants would increase the viral transmissibility and disease severity and reduce the protective ability of vaccines [10].

Besides the rapidly emerged SARS-CoV-2 variants, previous studies also reported the coinfection of SARS-CoV-2 with other respiratory pathogens [11, 12]. David Kim and his colleagues found that 116 COVID-19 patients were also positive for other microbial pathogens, such as influenza A/B, respiratory syncytial virus, human metapneumovirus, and *Chlamydia pneumoniae* [11]. Also, the reinfection with different SARS-CoV-2 variants in a COVID-19 patient has been reported. Richard L Tillett *et al*. presented a COVID-19 patient who tested positive for SARS-CoV-2 on April 2020 and was reinfected by a different SARS-CoV-2 variant on June 2020 [13]. The astonishing discovery was hard to explain why previous exposure to SARS-CoV-2 failed to provide immunity protection to the patient. Since coinfection is prevalent in viral infections [14-16], the studies inspire us to explore whether coinfection of multiple SARS-CoV-2 variants exists in COVID-19 patients, providing clues for prolonged viral shedding time and severe symptom [17].

Here, we collected 12,986 SARS-CoV-2 genomic sequences from the GISAID database on May 11, 2020, constructed single-nucleotide polymorphisms (SNP) coexistence network, and found a maximal clique of 16 coexisted loci. By simulating the SNVs accumulation with SARS-CoV-2 transmission, we discovered 2.20 averaged coinfected variants in the COVID-19 patients with the coinfection index. To validated the methods and results, we extracted 4,113 additional genomes from the GISAID database on April 1, 2021, and discovered an increased coinfected variants number of 3.42. Then, we performed Nanopore sequencing on the sputum samples from 42 COVID-19 patients and found the heterozygous SNPs on some loci of the SARS-CoV-2 genome, confirming the multiple variants coinfection. Hence, our study proposed a computational simulating method to detect the number of the coinfected variants in COVID-19 patients, confirmed the coinfection of multiple SARS-CoV-2 variants, and implied the increased coinfected variants in the epidemic.

## Materials and methods

### Ethics Statement

The First Affiliated Hospital approved this study of Guangzhou Medical University, and the sample and data collection procedures were conducted following the principles expressed in the Declaration of Helsinki. All patients provided written informed consent and volunteered to receive investigation for scientific research.

### GISAID datasets and mutation detection

This study collected SARS-CoV-2 genomic sequences from the GISAID database (https://www.gisaid.org/) and divided them into two genomic datasets according to their releasing date: For the 12,986 SARS-CoV-2 genomic sequences published before May 11, 2020, we noted them as GISAID20May11 dataset; For the 4,113 SARS-CoV-2 genomic sequences posted on April 1, 2021, we noted them as GISAID21Apr1 dataset. All genomes in these two datasets were tagged as complete (>29,000 bp) and high coverage (<1% Ns with <0.05% unique amino acid mutation) in the GISAID. We adopted MUMmer (version 3.23) to obtain the SNVs of the SARS-CoV-2 genomes [25]. Each SARS-CoV-2 genome is aligned with the SARS-CoV-2 reference genome (MN908947.3) to obtain the homology region using the nucmer function with the default parameters [25]. Then we got the SNPs matrix from the alignment results with show-SNPs function [25] and prepared for the SNV clique analysis.

### SNP coexistence network and clique analysis

To evaluate the complexity of SNPs co-occurrences within the GISAID dataset, we applied single-nucleotide variant (SNV) clique analysis by in-house scripts. Firstly, we considered a pair of SNPs from two different loci as complex if it occurred in at least one variant of the GISAID datasets. However, a complex paired-loci is hard to be explained in phylogeny, and it may happen by chance. Therefore, to remove such a possibility, we performed an analysis based on SNV cliques instead.

After obtaining all SNPs, we checked the alleles at every locus of the SARS-CoV-2 genome. Over 92% of the SNPs loci (5,671/6,178) had two alleles. Focusing on the loci with two alleles, we removed the SNPs loci with three or four alleles. We labeled the major allele of SNP locus as R and the minor allele as A. Thus, it had four possible genetic combinations for every pair of two SNPs loci: RR, RA, AR, AA. We recognized each SNP locus as a vertex and created an edge between a loci pair only if all four genetic combinations existed in at least one assembly genome within the GISAID dataset (Figure 1A). We obtained the maximal clique from the network. Based on the cliques, we can tell whether the SARS-CoV-2 coinfection exists since the existence of a large clique will be intractable to explain using phylogeny.

**Figure 1.**
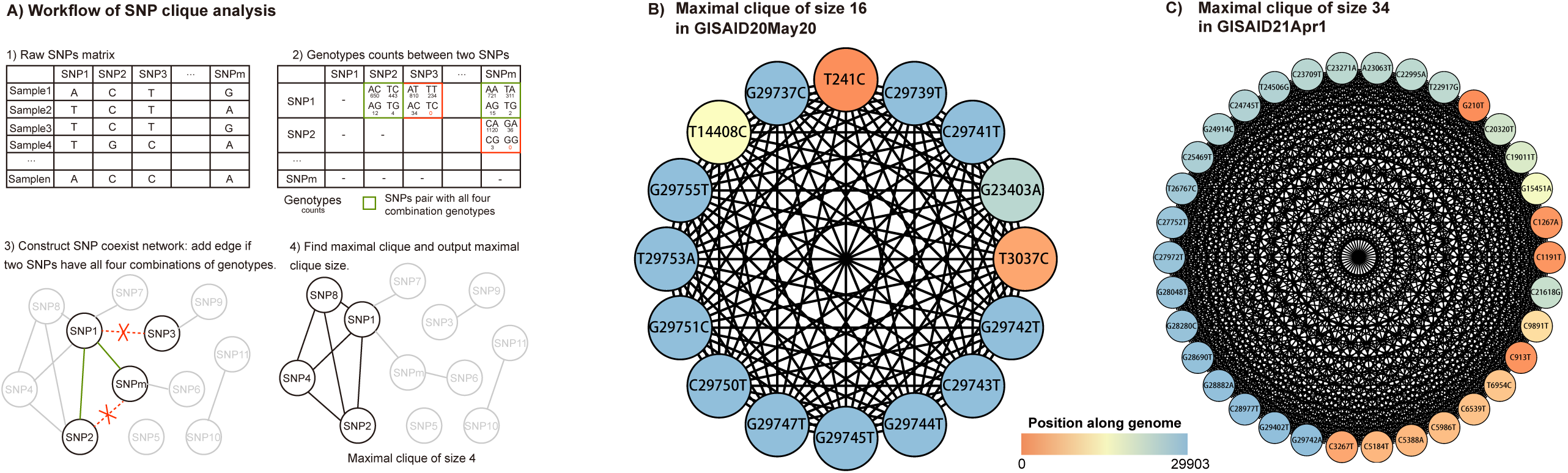
The workflow of SNP clique analysis and maximal clique in the collected GISAID datasets. **A)** First, we construct the SNP coexisted network from the SNP matrix. Every SNP locus is a vertex, and we add an edge between a loci pair if they have all four major genotypes. We then extract the maximal clique from the network. **B)** The maximal 16-SNV-clique was found in the GISAID20May11 dataset with 11,179 SARS-CoV-2 genomes. **C)** In the GISAID21Apr1 dataset, the 4,113 SARS-CoV-2 genomes contained the maximal clique of size 34.

### Prediction of coinfected variant number based on the simulation

With the SNVs in the collected genomic sequences, we predicted the coinfected variant numbers by simulations with the mutation rate (r) and the average variant number (w). In previous reports, the estimated mutation rate of SARS-CoV-2 by several groups ranged from 2.88×10-6 to 3.45×10-6 substitutions per site per day [26-28]. However, the obtained SNVs number distribution curve in our test does not fit the distribution curve from the real data set with a mutation rate of 3.0×10-6 (Supplementary Figure 1). The mutation rate we used has four values, which are 1.5×10-6, 2×10-6, 2.5×10-6, and 3×10-6. The average variant number in the simulation with 15 values ranged from 1.2 to 4, with an interval of 0.2. The distribution of variant numbers in all samples conformed to Poisson distribution with λ equals the average variant number.

### Sample collection

To confirm the coinfection of SARS-CoV-2 variants, we performed RT-PCR on the sputum samples collected from COVID-19 patients. Forty-two patients were recruited from the First Affiliated Hospital of Guangzhou Medical University and Guangdong Second Provincial General Hospital, China (Supplementary Table 1). The sputum samples from the patients were inactivated under 56°C for 30 minutes following WHO and Chinese guidelines [29-31]. The specimens were stored at 4°C until ready for shipment to the Guangdong Centers for Disease Control and Prevention.

### Nanopore sequencing

For the samples, we extracted the total RNA from the samples according to the protocol of RNA isolation kit (RNAqueous Total RNA isolation Kit, Invitrogen, China), and determined the RNA concentration by Qubit (ThermoFisher Scientific, China). Based on two pools of primers (98 pairs of primers in total) (Supplementary Table 2), the entire genomic sequence of SARS-CoV-2 was amplified segmentally by reverse transcription. Then, the libraries were built by adding the adapter and barcode to the amplified genomic fragments with a Nanopore library construction kit (EXP-FLP002-XL, Flow Cell Priming Kit XL, YILIMART, China). The samples were sequenced on the MinIon sequencing platform (Oxford Nanopore Technologies, U.K.).

### Nanopore data filtration

MinIon sequencer generated Fast5 format data, which was converted into fastq format with guppy basecaller (version 3.0.3). By applying NanoFilt (version 1.7.0) [32], we performed data filtration on the raw fastq data with the following criteria: the read lengths should be longer than 100 bp after removing the adapter sequences overall quality of reads should be higher than 10. Furthermore, due to the random connection of multiplex RT-PCR amplicons, the chimeric reads should be processed to avoid false identification of virus recombination or host integration. Therefore, we positioned the primers on the sequencing reads to identify the chimeric reads, split the identified chimeric reads into segments corresponding to PCR amplicons, and retained the final reads by aligning the segments to the viral genome (Supplementary Figure 2). This method allowed us to salvage a huge amount of sequencing data, leading to more accurate alignment and higher coverage.

### Mutation detection with Nanopore data

We aligned the filtered and segmented reads to the SARS-CoV-2 reference genome (MN908947.3) with Minimap2 by applying the default parameters for Oxford Nanopore reads [33]. The aligned PCR amplicons were separated according to the corresponding primer pool. With the separated alignment results, the genomic variations with average quality larger than ten were called with bcftools (version 1.8) [34]. Mutations with less than ten supported reads were filtered. To reduce the PCR amplification effects, we also filtered the variations within ten bp upstream or downstream of the primer region within the corresponding primer pool. The filtered mutations for different primer pools were then merged as the final mutations. The final mutations were annotated by in-house software based on the gene information in the SARS-CoV-2 reference genome.

## Results

### Discovery of the 16-SNV-clique with the GISAID20May11 dataset

The GISAID20May11 dataset contains 12,986 SARS-CoV-2 genomes published between December 30, 2019, and May 11, 2020. After filtering 1,804 duplicated sequences, we aligned the rest of 11,182 viral genomes to the SARS-CoV-2 reference genome to obtain SNVs. Then, we removed three viral genomes with over 1,000 SNVs and obtained 11,179 genomes for the following-up analysis. With 57,548 SNVs on 6,178 SNPs loci, we performed SNP clique analysis (**Figure 1A**) and constructed the SNP coexistence networks with 1,150 vertices and 8,003 edges. Among the networks, we discovered the maximal clique with 16 coexisted loci (**Figure 1B**). With the result, we deduced that some SARS-CoV-2 assembly genomes were mixed sequences of multiple coinfected variants, except the incredible-fast mutation.

### Coinfection index to determine the SARS-CoV-2 variant number in a sample

We selected the maximal clique from the SNP coexistence networks and noted its size as the coinfection index. We further determine the average coinfected variants number with computational simulations. By simulating the transmission route tree of COVID-19, we traced the virus transmission among the infected individuals. Based on the publishing date of the sequences, we selected the sequences at the same transmission period as the simulated sequences and calculated the coinfection index using SNP clique analysis. Using different mutation rates and the average variant number in the simulation, we could obtain a chart of the average variant number against the coinfection index under a specific mutation rate (**Figure 2A**). During transmission, the variants in a sample at the child node were randomly inherited from the sample at the parent node. The variants would generate new SNVs based on a given simulated mutation rate (**Figure 2B**). In the simulation, we proposed two methods of how the coinfected variants in a sample construct their assembly genome. The first method randomly selected a variant from the coinfected multiple variants in the sample, and reported the SNVs in this variant. The second method (the mixed method) generated an assembly genome, which was a mixture of all variants. We split the genome as windows with a fixed size of 100 bp for the second method, and each window comes from a randomly selected variant in the sample. Using these two methods, we obtained the SNVs in the assembly genome (**Figure 2C**).

**Figure 2.**
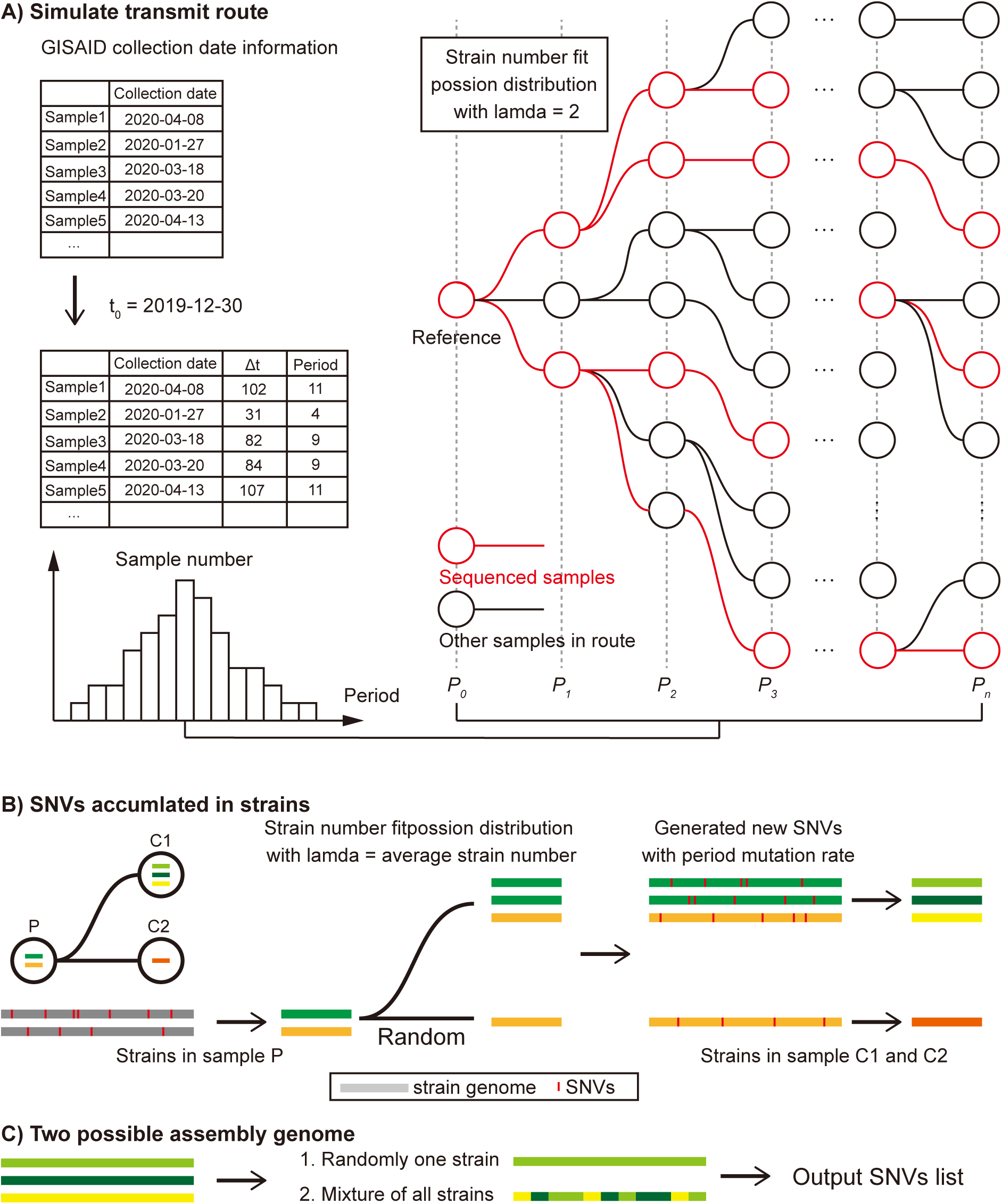
The simulation flowchart of viral SNVs in samples. **A)** We simulated the transmitted route based on known epidemiological information of SARS-CoV-2, and construct the transmission tree. Then we select the sequenced samples based on their releasing date in GISAID database. **B)** Variants number in all samples fit the Poisson distribution with λ equals the average variant number. In a single transmission branch, variants in child nodes are randomly inherited from the parent sample. For every child variant, we generated new SNVs with the period mutation rate. **C)** We simulated two possible assembling situations of samples with multiple variants coinfection and acquired the SNVs list of all samples as the output.

After plotting the coinfection index against the average variant number, we got two regression lines between them (**Figure 3A**). With the results, we noticed that only the regression line based on the mixed method could achieve a coinfection index of 16 for the GISAID20May11 dataset. With the regression lines of these two methods, we concluded that the 16-SNV clique from the GISAID20May11 dataset should result from coinfection, and the assembly genome comes from the mixed sequencing data of the coinfected variants.

**Figure 3.**
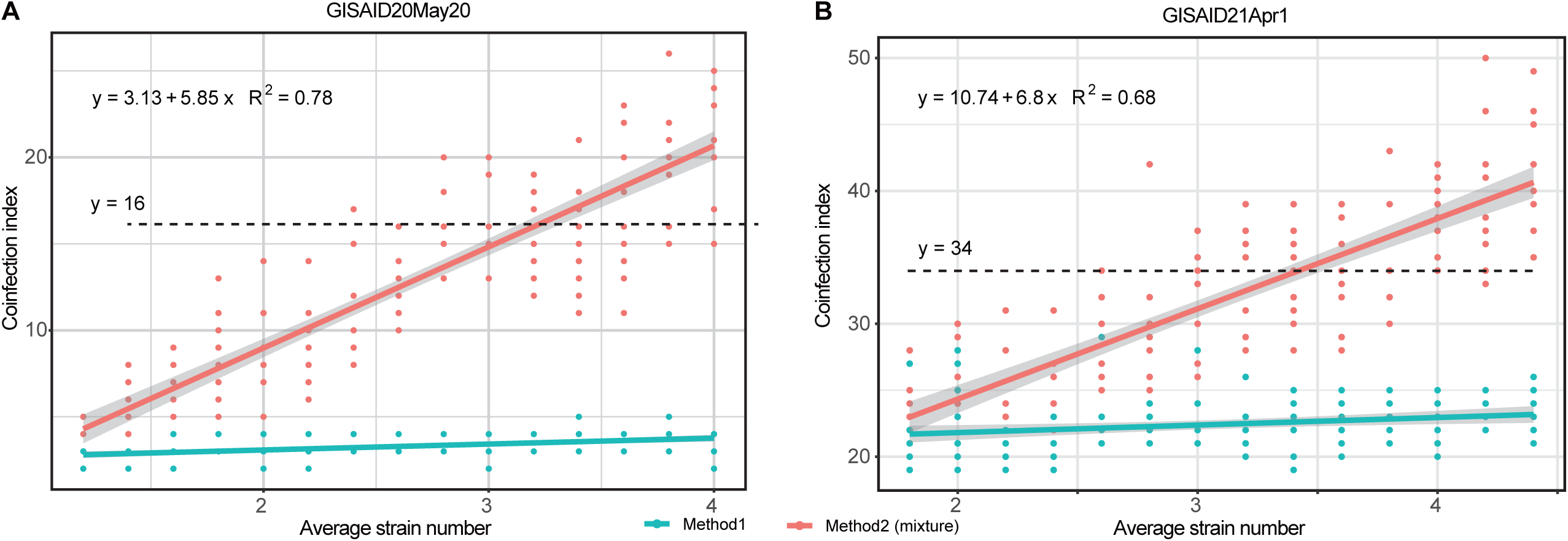
The regression of variant number and the coinfection index. **A)** The distribution of coinfection index with different average variant numbers in the GISAID20May11 dataset. Method 2 exhibited the linear regression relationship between the coinfection index and average variant number, and the generated formula suggested the mixed variants of the assembly genome in the dataset. **B)** The linear relationship between coinfection index with different average variant numbers in the GISAID21Apr1 dataset. With method 2, the average variant number was 3.4 when the coinfection index was 34.

Then, we determined the averaged variant number in the GISAID20May11 dataset with the coinfection index line. We performed regression analysis between averaged variant number and coinfection index and discovered the significant linear relationship between them with method 2 (F-statistic p-value < 2.2e-16, adjusted R-squared = 0.79, Figure 3A). According to the obtained fitting equation, we deduced that the corresponding average variant number was 2.20 when the coinfection index was 16 (Figure 3A).

### Coinfection index increased along with the COVID-19 pandemic

With the GISAID20May20 dataset, we obtained a maximal clique with 34 coexisted SNPs from 140,348 SNVs on 6,415 SNPs loci (**Figure 1C**). Then, we constructed the coinfection index curve with the GISAID21Apr1 dataset and determined the average variants number in this dataset. The genomes of GISAID21Apr1 were sampled from five different continents. Europe provided primary samples as 3,023 samples were from Europe, and the rest 1,047 samples were from North America, 27, 12, and 4 samples were from Asia, South America, and Oceania, respectively. While, we found 28 SNPs existed in over 3,000 samples, which reveals those samples should have the same or related ancestor. We altered the simulated procedure since we assumed those samples had the same ancestor to fit the SNVs distribution in the GISAID21Apr1 dataset. The regressed linear of the coinfection index and the average number of variants showed a significant linear relationship (F-statistic p-value < 2.2e-16, adjusted R-squared = 0.69, **Figure 3B**). The fitting equation revealed the average stain number of 3.42 in the GISAID21Apr1 dataset. The pandemic of COVID-19 made the virus could transfer between continents and increased the coinfection of different variants.

### Sequencing data statistics for the 42 COVID-19 patients

For the 42 COVID-19 patients enrolled from the First Affiliated Hospital of Guangzhou Medical University and Guangdong Second Provincial General Hospital, we performed Nanopore sequencing on their sputum samples for SARS-CoV-2 genome acquirement and mutation detection (Supplementary Table 1). After sequencing on the multiple-PCR products, a total of 7,877,736 clean reads were generated, with an average of 187,565±143,719.55 (Mean±SD) reads per sample (**Figure 4**). To eliminate the chimeric reads formed by the unintended random connection of multiplex PCR amplicons, we developed a software tool named CovProfile [18] (Supplementary Figure 2) and perform data filtration and detect the mutations in SARS-CoV-2 variants. Then we discovered that the chimeric reads were making up 1.69% of total sequencing reads. Aligning the clean reads to the SARS-CoV-2 genome and human transcriptome, we discovered that the ratio of primary aligned sequence ranged from 3.86% to 99.74% on the SARS-CoV-2 genome and ranged from 0.13% to 70.5% on the human transcriptome database (Figure 4). Moreover, the SARS-CoV-2 genomic coverage reached over 99.7% with >1800x depth in each sample, ensuring adequate data volume for SNP calling (Supplementary Figure 3).

**Figure 4.**
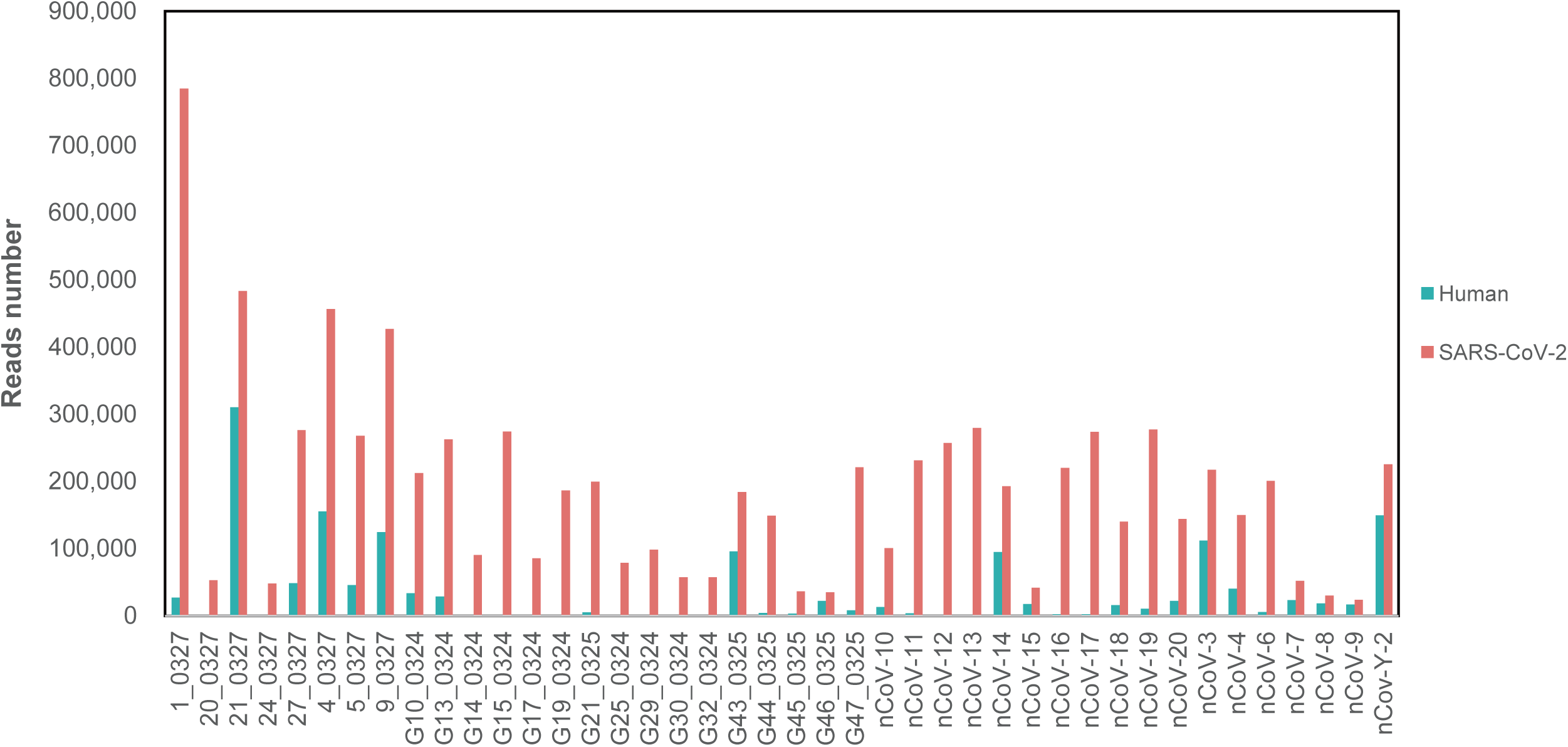
Statistics of Nanopore sequencing data for the 42 COVID-19 samples. After the low-quality filtration, we aligned the sequencing data to the SARS-CoV-2 genome and human transcriptome, respectively. The histograms in red and green represent the reads number aligned to the SARS-CoV-2 genome and human transcriptome.

### Identification of heterozygous SNPs on SARS-CoV-2 genome

After aligning the filtered data to the SARS-CoV-2 genome, we detected the mutations of SARS-CoV-2 in the 42 samples (**Figure 5**). Based on these mutations, we discovered a total of 115 SNPs in all samples, and 108 of them located on the genetic regions, including genes ORF1ab, S, ORF3a, N, M, ORF6, ORF8, and ORF10 (Supplementary Table 3). Furthermore, we discovered the heterozygous SNPs in 41 of the enrolled samples (Figure 5). Since each locus contained only one genotype in a viral genome, the heterozygous SNPs indicated that each host was infected with two variants at least. Moreover, twenty heterozygous SNPs existed in over two samples, such as C865T, A1430T, C8782T, etc (Supplementary Table 3). Notably, we also discovered that 14 samples contained two genotyped SNPs on loci 8,782 and 28,144 simultaneously, which were significant SNPs identified in recent phylogenetic analysis. Meanwhile, we did not find creditable InDels (Insertions and Deletions), structural variations, or viral-host recombination.

**Figure 5.**
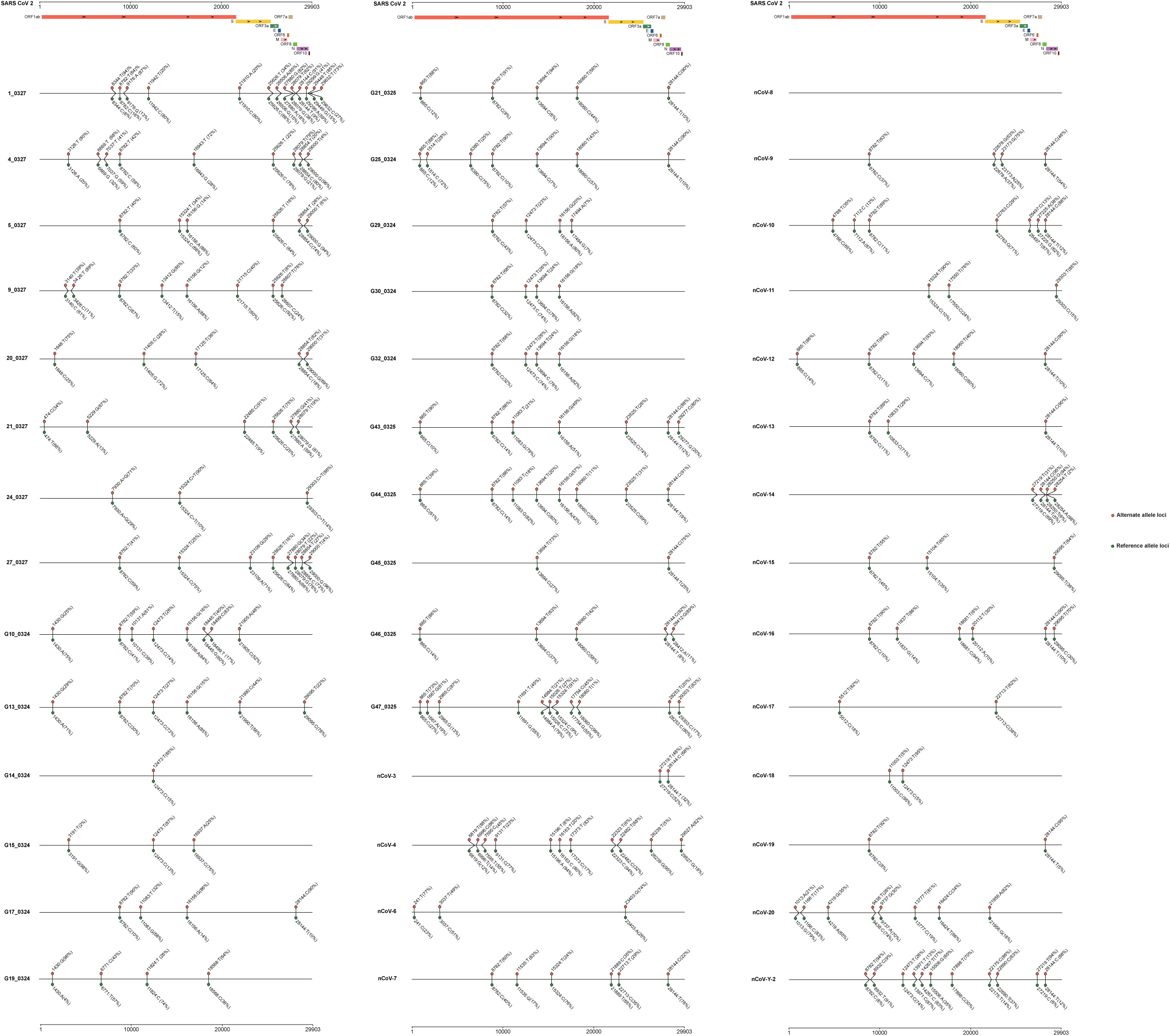
SNP distributions in 42 samples gathered from COVID-19 patients. The alternate alleles were shown in red, while the reference and mutated alleles were in green and red, respectively.

## Discussion

SARS-CoV-2 posed a significant threat to human lives, and recent studies have reported the rapidly emerged variants and their impact on clinical severity and vaccine protection [7, 9, 19]. In this study, we aimed to detect whether the coinfection of multiple SARS-CoV-2 variants exists in COVID-19 patients, which might associate with frequent homologous recombination and greater clinical severity. This study performed the SNP coexistence network analysis to detect the “coinfection index” based on the maximal clique in the collected GISAID datasets and constructed the relationship between the coinfection index and the average variant number. We deciphered the number of coinfected variants for SARS-CoV-2 in hosts with the linear regression between the coinfection index and the average variant number. With the GISAID20May20 and GISAID21Apr1 datasets, we discovered that the number of the coinfected variants increased from 2.20 to 3.42 in the COVID-19 patients. Considering the rapidly emerged SARS-CoV-2 variants worldwide, we hypothesized that the coinfected variants in hosts would aggravate the clinical severity, increase the change of viral recombination, and posed a greater threat to us [20]. Moreover, the coinfection index can be applied to other viruses in hosts. Although the coinfection explained the large clique detected in the SNP coexistence networks in the collected datasets, the discoveries still need to be verified experimentally.

To verify the coinfection of multiple SARS-CoV-2 variants, we performed Nanopore sequencing on 42 COVID-19 patients and implemented CovProfile for the sequencing data processing and the genomic mutation detection [18]. Our results confirmed the reliability of the multiplex RT-PCR method in identifying SARS-CoV-2 and discovered the recurrent heterozygous SNPs on 41 of 42 samples. Moreover, we found two genotyped SNPs on loci 8,782 and 28,144 in fourteen patients. Since loci 8,782 and 28,144 were important for SARS-CoV-2 phylogenetic analysis [21], the finding has crucial impacts on the evolution derivation of SARS-CoV-2, as the heterogeneous loci might cause mis-links during viral genomic assembly. Corresponding to the simulation results, the discoveries of heterozygous SNPs confirmed the multiple variants coinfection in the COVID-19 patients.

The discovery of SRAS-CoV-2 variants coinfection provided explanations for the severe clinical symptoms in some COVID-19 patients and significantly impacted the application of vaccines [9, 22, 23]. Since vaccines were developed referencing a specific SARS-CoV-2 variant, the infection of variants limited the protection afforded by vaccines [9]. For instance, SARS-CoV-2 B.1.351 variant, which is widely spread in Nelson Mandela Bay and South Africa, can evade the immune response stimulated by the vaccines and greatly reduce the vaccine’s protective effect on the population [19]. Moreover, Nicole Pedro et al. also discovered the coinfection of dual SARS-CoV-2 variants in a severity COVID-19 patient in Portugal, which supported our discoveries [17]. Therefore, the coinfection of multiple SARS-CoV-2 variants raised another challenge, and we need to stay alert in the battle against the COVID-19 epidemic.

Although the findings implied the coinfection of multiple SARS-CoV-2 variants in patients from the perspectives of algorithm derivation and mutation detection, this study still has several limitations. In the simulation, we assumed that the first submitted sequence was the source of all SARS-COV-2 variants. While, in the pandemic, the first infective SARS-COV-2 variant should emerge long before being discovered. The study by Giovanni Apolone *et al*. proposed that SARS-CoV-2 RBD-specific antibodies can already be detected in the serum samples of Italian cohorts collected in March 2019, indicating that the source variants of all currently sequenced variants should appear earlier before [24]. Determining the virus’s origin is difficult, so we chose an exact time point during the simulation, but it does not affect our conclusions on the coinfection of multiple variants in hosts. Moreover, there was no guarantee considering the quality of the viral variants submitted to GISAID, which might influence the accuracy and potential phylogenetic study. Last but not least, the discovered heterozygous SNPs need to be verified with biological duplication, and we should identify the coinfected viral lineages in the COVID-19 patients in future study.

In conclusion, our study proposed a computational simulating approach to decipher the number of the coinfected variants, declared the coinfection of multiple SARS-CoV-2 variants in COVID-19 patients, and reported the increased coinfected variants in the COVID-19 epidemic, reminding us of the threats brought by the SARS-CoV-2 infection.

## Data availability

CovProfile is an open-source collaborative initiative available in the GitHub repository (https://gitlab.deepomics.org/yyh/covprofile). All other code is available from the authors upon reasonable request. The Nanopore sequencing data in this paper have been deposited in the Genome Sequence Archive in BIG Data Center, Beijing Institute of Genomics (BIG), Chinese Academy of Sciences, under BioProject PRJCA002503 with accession ID CRA002522 (https://bigd.big.ac.cn/gsa).

## Authors’ contributions

S.C.L., B.S. and F.Y. proposed the simulation approach and supervised the project. Y.L. and Z.L. performed the samples collection and Nanopore data analysis. Y.J. and Y.Y. collected the public data and optimized the algorithms in simulation. J.C., W.J. and Y.K.N. guided the analysis and optimized the graphs. Y.L., Y.J., Z.L. and Y.Y. interpreted the results and wrote the manuscript. S.C.L., B.S. and F.Y. polished the manuscript. All authors reviewed the article and approved the final manuscript.

## Acknowledgments

This work was supported by the Chengdu Science and Technology Project for COVID-19 prevention and control [Grant no. 2020-YF05-00281-SN] awarded to B.S. We would like to thank Beijing YuanShengKangTai (ProtoDNA) Genetech Co. Ltd. for their helping in Nanopore sequencing, and we would also like to thank all the doctors in the First Affiliated Hospital of Guangzhou Medical University, for their assistance on the specimen and clinical data collection.

## Supplementary material

**Supplementary Figure 1. The distribution of samples with different SNVs in GISAID20May11 dataset and the simulation under different mutation rates**.

We had 15 possible average numbers of variants and ten duplicates for each pair of mutation rate and the average variant number. We plotted sample number in all simulations and regress samples number VS number of SNVs of all simulations with specific mutation rate, and the 95% CI region showed in grey.

**Supplementary Figure 2. The procedure of chimeric reads identification and reads splicing**.

**Supplementary Figure 3. The coverage of depth of aligned data in the 42 COVID-19 samples**.

The X coordinate stands for the location of SARS-CoV-2 genome, and the Y coordinate stands for the sequencing depth. The bars with red, yellow, green, pink, brown, light green, purple and dark brown colors stand for the genetic regions of ORF1ab, S, ORF3a, M, ORF6, ORF8, N and ORF10, respectively.

**Supplementary Table 1. Physical information of the 42 enrolled COVID-19 patients**.

**Supplementary Table 2. Primers applied for RT-PCR amplification of SARS-CoV-2**.

**Supplementary Table 3. SNP distributions on 42 COVID-19 patients**.

